# Hidden patterns of codon usage bias across kingdoms

**DOI:** 10.1101/478016

**Authors:** Yun Deng, Fabio de Lima Hedayioglu, Jeremie Kalfon, Dominique Chu, Tobias von der Haar

**Affiliations:** School of Computing, University of Kent, CT2 7NF, Canterbury, UK; Kent Fungal Group, School of Biosciences, University of Kent, CT2 7NJ, Canterbury, UK; Broad Institute of Harvard and MIT, Cambridge, MA 02142, Cambridge, USA

**Keywords:** stochastic thermodynamics, codon usage bias, fungi, protists, bacteria

## Abstract

The genetic code is necessarily degenerate with 64 possible nucleotide triplets being translated into 20 amino acids. 18 out of the 20 amino acids are encoded by multiple synonymous codons. While synonymous codons are clearly equivalent in terms of the information they carry, it is now well established that they are used in a biased fashion. There is currently no consensus as to the origin of this bias. Drawing on ideas from stochastic thermodynamics we derive from first principles a mathematical model describing the statistics of codon usage bias. We show that the model accurately describes the distribution of codon usage bias of genomes in the fungal and bacterial kingdoms. Based on it, we derive a new computational measure of codon usage bias — the distance 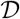 capturing two aspects of codon usage bias: (*i*) Differences in the genome-wide frequency of codons and (*ii*) apparent non-random distributions of codons across mRNAs. By means of large scale computational analysis of over 900 species across 2 kingdoms of life, we demonstrate that our measure provides novel biological insights. Specifically, we show that while codon usage bias is clearly based on heritable traits and closely related species show similar degrees of bias, there is considerable variation in the magnitude of 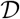 within taxonomic classes suggesting that the contribution of sequence-level selection to codon bias varies substantially within relatively confined taxonomic groups. Interestingly, commonly used model organisms are near the median for values of 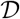 for their taxonomic class, suggesting that they may not be good representative models for species with more extreme 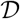, which comprise organisms of medical an agricultural interest. We also demonstrate that amino acid specific patterns of codon usage are themselves quite variable between branches of the tree of life, and that some of this variability correlates with organismal tRNA content.

## Introduction

Codon Usage Bias (CUB), the preferential use of some types of codons over others encoding the same amino acid during protein synthesis, is now an empirically well established phenomenon. Yet, it remains unclear why CUB has evolved and which evolutionary forces drive its evolution. Existing approaches to studying codon usage use a number of measures for the frequency of individual codons relative to particular, “optimal” reference codons. Such measures include the frequency of optimal codons *F_opt_* [1], the codon bias index *CBI* [2], and the codon adaptation index *CAI* [3]. These heuristic measures can be usefully contrasted to model-based approaches, where it is assumed that codon usage is shaped by a force or forces that affect the distribution of codons. Existing modelbased approaches include the “Effective Number of Codons” (*EN_c_*) which essentially performs a statistical test against the null hypothesis that codon usage is solely governed by genomic GC content [4], models based on the combined forces of mutation bias and selection for minimal energy usage during translation [5, 6], and models based around the forces of mutation, selection and genetic drift in populations [7, 8]. Both heuristic and model based approaches rely on a number of assumptions as well as contextual knowledge about the organisms under study. For example, measures of codon optimality require defining sets of highly expressed genes, whereas model based approaches typically require knowledge of specific molecular and evolutionary mechanisms. Moreover, current model-based approaches limit their scope to a small number of forces shaping CUB, even though many more forces have been proposed to affect CUB in the literature.

In general terms, one can think of two classes of forces acting on codon usage. (*i*) Selection forces may directly shape the genome-wide relative frequency of codons, in a manner that affects all genes uniformly. We will henceforth refer to this as *beanbag selection*, because codon choice would then be analogous to selecting different coloured beans from a bag containing these colours in a given proportion. Mathematically, beanbag selection-type forces would lead to a CUB where the genome-wide codon distribution is biased and codon usage is distributed across genes according to a multinomial distribution. (*ii*) Alternatively, it is conceivable that selection operates on individual genes, for example such that some combinations of codon choices are preferred for some genes, whereas other combinations are preferred for other genes. We will refer to this as *sequence level selection* (SLS). This, in contrast to beanbag selection, will yield a non-multinomial distribution, and genome-wide codon frequencies that may or may not appear biased.

There are plausible, well characterised mechanisms for both beanbag selection and SLS. Beanbag selection includes mechanisms such as mutation biases [5, 6, 9], biases in genomic GC content [4], genetic drift [10], and global selection for efficient codons to optimise ribosome usage [11]. SLS includes selection for codon usage that is compliant with particular expression levels (for example, for efficiently decoded codons in highly expressed genes [12], or for inefficiently decoded codons in genes where low expression level is important [13]), and selection against translational errors at structurally sensitive sites in proteins [14]. It is worth noting that this distinction between SLS and beanbag selection is idealised, and real biological forces may have components of both. For example, avoidance of translational errors may well shape CUB genome wide, but may be particularly important at structurally sensitive sites in proteins. The literature also continues to propose additional forces that may shape CUB [15–20], which are not well enough understood to clearly assign them as either beanbag selection or SLS. Nevertheless, the central point that different types of forces exist which have very different consequences for CUB remains valid.

With the explosion in genome sequencing data, there is an emerging scope for studying CUB and sequence evolution at a scale much broader than ever before. However, for organisms where data about tRNA populations and other parameters is not available — and this is the case for the vast majority of genome sequences — applying the established approaches can be problematic. For example, current methods cannot be applied to many interesting organisms, such as organisms with extreme lifestyles like parasites [21–23] or thermophiles.

In this contribution, we shall propose a fresh approach to quantifying CUB that does not rely on a choice of comparison sets, nor does it assume a particular mechanistic model of codon usage bias. As a consequence it can be applied to the wealth of emerging genomic information, including genomes from poorly characterised organisms. Drawing on ideas from statistical physics and stochastic processes, we develop a novel quantification method — the *distance measure* 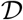 — that expresses the amount of bias in a particular genome. The significance of this measure is that it encapsulates both the bias arising from beanbag selection, as well as bias arising from SLS. Furthermore, the calculation of 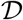 does not rely on any *a priori* choice of gene reference sets. The development of 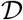 is the first main contribution of this article.

As a second main contribution we then use 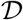 to analyse ≈ 1500 genomes from across 3 microbial kingdoms of life. This analysis will show that (*i*) 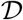 captures relevant biology, (*ii*) SLS strongly contributes to shaping overall codon usage bias in most organisms, (*iii*) amino acid specific patterns of codon usage are variable between branches of the tree of life, and (*iv*) that 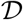 varies with the ecological niche of the organism.

## Results

### Codon selection as a random walk

We start by deriving a probability distribution for codon usage. In order to do so, we conceptualise codon evolution as a discrete space, continuous time random walk in the space of synonymous codons. In this picture, each gene represents up to 18 independent random walkers, one for each of the amino acids that is encoded by more than 1 codon. For the purpose of this article, we will exclusively focus on synonymous mutations, whereby a codon *a* encoding an amino acid gets exchanged for a codon *a*′ that encodes the same amino-acid. We can then conceptualise each gene sequence as consisting of (up to) 18 independent subsequences, where each subsequence is a string of codons encoding the same amino acid 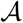 appearing in a gene *g* and 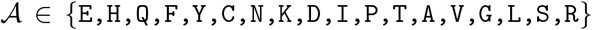. For each of these subsequences, we then counted how often of each of the synonymous codons appears. In this way, we obtained what we will henceforth call the *observed codon count* (OCC); see fig. 1 for a graphical explanation of the OCC.

**Fig. 1:**
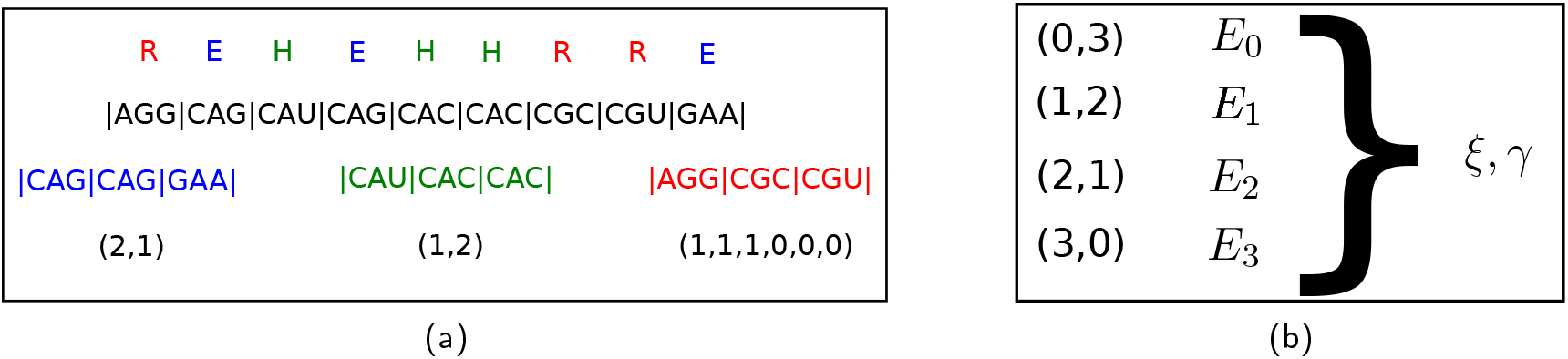
(a) For each species, each mRNA is split into up to 18 OCCs, i.e. one OCC for each amino acid with a codon choice greater than one. Here, we show the example of a hypothetical mRNA coding for `REHEHHRRRE`. This mRNA is split in three subsequences, one for each of its constituent amino acids. The third line shows the subsequences of codons split up by amino acid in the order `E,H,R`; see color code. The last line shows the OCCs for this gene, i.e. the counts of the codons. In this case, all OCCs happen to be of length *L* = 3. A synonymous mutation could take an amino acid from state, say (2,1) to either (3, 0) or (1, 2). The “energy” of an OCC is then calculated based on the number of codons of each type in each OCC, as given in the last line. (b) For each amino acid of an organism, the OCCs for each length *L* are collected; here an example is shown for *L* = 3. For each configuration the frequency of configurations is calculated and fitted to a Boltzmann distribution based on eq. 6 to obtain *ξ* and *γ*. This in turn is used to calculate the distance 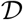 according to the formula.

For our random walk model, we will consider each of the OCCs as an independent random walker. Throughout this manuscript, we denote the number of codons for amino acid 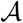 in gene g by 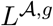. Each OCC consists of *k*_1_ codons of type 1, *k_2_* codons of type 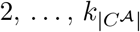 codons of type 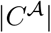, where 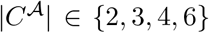 is the total number of codons for amino acid 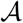. For each amino acid, we arbitrarily assigned codons to codon 1, codon 2 and so on. This assignment remained fixed for all species we analysed. Table 1 summarises the symbols used.

**Tab. 1:**
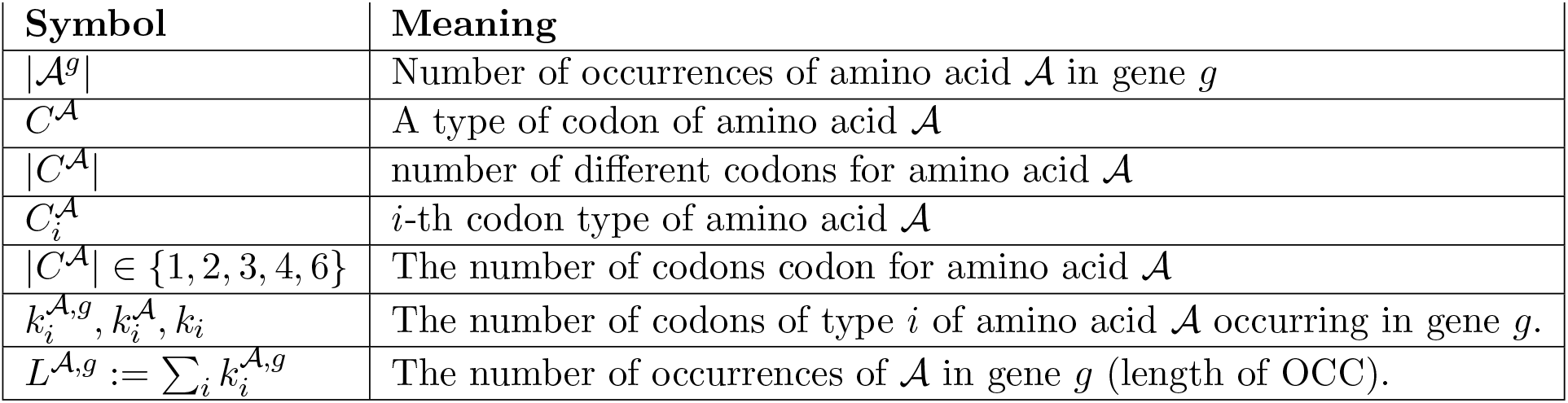
Explanation of the symbols used.

We can now consider each possible OCC 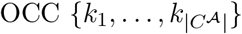 (of a particular amino acid) as a state of a random walker; see fig. 1 for a graphical explanation of what we mean by “state.” From any such state, the random walker can access all states that are 1 synonymous mutation away. For example, one of the codons in the OCC may be mutated from codon 1 to codon 2, which would correspond to the transition from 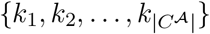 to 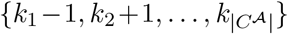. In the case of only two codons, where 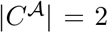 this random walk reduces to a 1-dimensional discrete state random walk in continuous time with 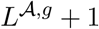 states; see SI sec. 1 for more detail.

Throughout this contribution, we make a number of simplifying assumptions about the nature of the random walk. Firstly, we do not consider non-synonymous mutations, i.e. the rate of mutation from a codon to a non-synonymous codon is zero. Secondly, we assume that the mutation rates between synonymous codons are *a priori* the same, i.e. the random walk is unbiased. Any deviations from this assumption arise from biasing effects that operate on the genome. These could be, for example, mutation biases, GC-bias or evolutionary selection pressures (including effects of random drift). A subtlety arises here in the context of 6 synonymous codons where the mutation rates between synonymous codons are not equal. This does not affect the discussion here, however, because we will focus mostly on amino acids with 2 synonymous codons. Thirdly, we assume throughout that the random walker is in a steady state. Continuing evolutionary pressure could therefore change individual OCCs, but will not, on the whole, change the statistics of the codon distribution. Fourthly, throughout this article we are not concerned with the spatial arrangements of codons across a gene or genome, but we only record the OCCs, i.e. the count of synonymous codons of a gene separated into amino acids.

### Deriving the full model

To derive predictions for the distribution of codons across OCCs in response to specific selective pressures, we devised a theoretical model of the dynamics of codon evolution based on stochastic thermodynamics [24]. Above we noted that each OCC *i* can be regarded as a state of a random walk. The synonymous mutation rates are the transition rates of the random walk.

To make some progress, we will now take this conceptualisation of a random walk seriously and imagine that the OCC random walker is in fact an abstract thermal particle that performs random transitions between some physical states. The transition rates between the states would then be due to energy differences between the states, such that each state i would then be associated with an energy *E_i_*. In the context of physical systems, the transition rates and the energies are related by the local detailed balance condition [25]

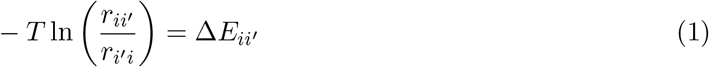

Here *r_ii′_* are the transition rates of the random walker to transition between a state with energy *E_i_* and *E_i′_*, with Δ*E_ii′_*: = *E_i_* – *E_ii′_*. Note that *r_ii′_* will be different for different pairs of *i* and *i′*. This equation allows us to calculate energies of individual states, if the rates *r_ii′_* are known, provided that one assigns an arbitrary 0 energy to some chosen state. The energies can then be used to calculate the associated Boltzmann (probability) distribution.

Returning to the random walk of the OCCs, we can now use this random walk conceptualisation in order to calculate the expected codon distributions under various assumptions about the transition rates. The simplest energy function can be derived for the beanbag model and in the special case of no selection forces acting on codon usage. The assumption of an un-biased codon selection implies that each codon has a constant mutation rate. Such a system is well known to generate a binomial distribution with *p* = 1/2 in the case of 2 codons, or a multinomial distribution with 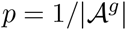 in the case of more than 2 codons.

It is instructive to show that using the physical interpretation of our random walk model, we consistently obtain the binomial distribution. In the unbiased case, the transition rates between OCCs are proportional to the number of codons, thus yielding the following random walk:

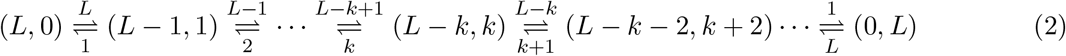

Using the local detailed balance condition (eq. 1) and because we choose the energy associated with state (0) to be *E*_0_: = 0, the energy difference between state (0) and state (1) is Δ*E*_1_ = – ln(*L*/1). Generally, the energy difference Δ*E_i_* between state (*i*) and state (*i* − 1) is given by Δ*E_i_* = −ln((*L* − *i* + 1)/*i*). Consequently, the energy of state (*k*) is then:

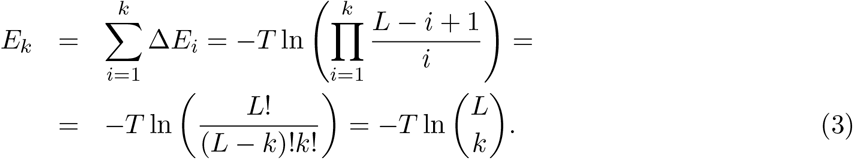

This entails that the (Boltzmann-)probability:

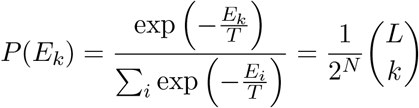

For *T* = 1 this is the binomial distribution with *p* = 1/2, as expected.

This simplest model can be readily expanded to include a global codon usage *bias q*, yielding an energy *Ê_i_* = *E_i_* + ln ((1 − *q*)^*i*^/*q^i^*) (for the calculation see SI sec. 1). The resulting Boltzmann distribution coincides again with the binomial distribution, but this time with a bias *p* = *q* ≠ 1/2.

The two preceding scenarios represent beanbag models, because they assume that the rate of mutation from codon 1 to codon 2 is proportional to *k*, the number of codons of type 1. We now posit instead that this rate is proportional to *k^ξ^* where 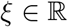, and the rate from codon 2 to codon 1 becomes proportional to 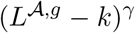 and 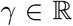. The resulting random walk model is then one where the rate with which codon 2 mutates to codon 1 and *vice versa* is no longer proportional to the number *N* − *k* + 1 and *k* of codon 2, but proportional to a power of this number. This leads to the following random walk model:

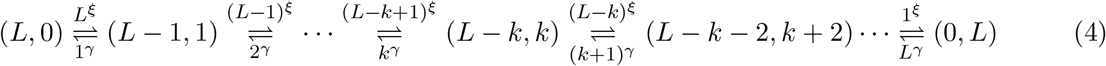

The rates of this random walk break the assumptions of beanbag selection in that the resulting statistics can no longer be reproduced by throwing dice or tossing coins, not even unfair ones. Following the same procedure as above, we can again calculate the energies for each configuration.

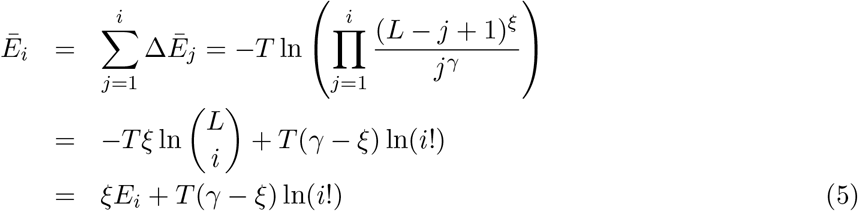

We thus obtain the *full model* of the distribution of codon usage.

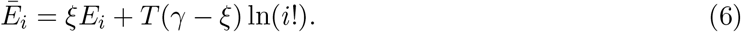

We can now associate an energy 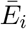 with each site *i* of the random walk. By construction each site corresponds to a specific OCC {*k*, *L* − *k*}_*i*_, thus mapping energies to OCCs. Altogether, this enables us to write down the steady state probability of observing an OCC as a Boltzmann distribution.

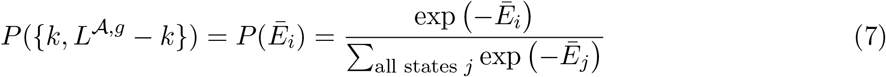

The full model eq. 6 and the associated Boltzmann distribution eq. 7 will form the core part of the analysis to follow. A few remarks on the model and the meaning of its parameters are useful at this point. The expression of the energy in the full model has two parts that lend themselves to direct interpretation. The first term on the left hand side is an “entropic” part that characterises codon usage in a no-selection scenario. This means that it does not lead to a global codon usage bias by itself. The second term is an effective “selection potential.” It encapsulates forces on the genome that alter the overall frequency of codons and thus modifies the probability distributions of the random walkers relative to the purely entropic case of no selection. We do not claim that this potential has a concrete single counterpart in biology. Instead, we interpret it as the emergent result of the variety of evolutionary forces that act on the genome.

The biological meaning of the ad-hoc parameters *ξ*, *γ* can be elucidated by considering some special choices for their values. When *ξ* = *γ* ≠ 1, then the second term on the right hand side disappears and the energy is the same as in the unbiased beanbag model with a modified inverse temperature *ξ*. In this case, there will be no selection pressure on the global usage of codons, but there may be SLS, affecting how codons are distributed across OCCs. For *ξ* = *γ* = 1 the full model eq. 6 reduces to the binomial distribution with *q* = 0.5 exactly. In the most general case of *ξ* ≠ *γ* selection is affected by the second term, which can be interpreted as a selection “potential.” In this case, a global codon frequency bias *q* will emerge as a result of SLS.

We note that the full model eq. 6 does not reduce exactly to the binomial distribution for *q* ≠ 1/2 for any choice of parameters, but we found that it can approximate it to high degrees of accuracy. Given the relatively high statistical error of determining codon distributions it will therefore not be possible to reject SLS empirically even if the underlying data was truly binomial.

Taking advantage of the fact that the *γ* = *ξ* = 1 entails the case of a sequence without any codon usage bias, it is possible to define, for each genome, a measure of how far it is away from the case of no codon usage bias. To do this, we use the (Euclidean) *distance* of a genome in *ξ*, *γ* space from a (hypothetical) unbiased, entirely random sequence located at *ξ* = *γ* = 1.

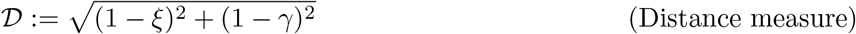

A vanishing distance 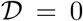 would indicate that no codon usage bias whatsoever is present in the genome as a whole. In this case, individual genes may still have measurable biases in codon usage but the biases would be distributed according to the binomial distribution, with no overall codon preference. When 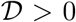 then this indicates that a CUB is present. The distance measure captures two types of biases: (*i*) a statistical global over-representation of the codon frequencies, and (*ii*) a deviation from the beanbag assumption. Either of those would be sufficient to lead to a non-vanishing 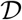. As such 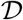 is useful as a high level descriptor of codon usage that provides a quantification of CUB complementary to comparable existing measures like *F_opt_* or *CAI*. Its value over and above existing measures is that it compares how far the distribution of codons is different from what would be expected in the case of no bias. As such, it could detect a bias in the case where the overall mean usage of codons is equal, but codons are distributed non-randomly across genes. We will see below that such cases exist. We will also show that 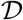 increases with the diversity of forces that shape codon usage in a set of sequences, such as mutation bias, GC content, or selection. Finally, a further feature of 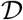 is that it does not rely on the choice of a reference set of any kind, but it can be computed directly from the genome data.

### Genomic data bear the signature of SLS

We now apply the model to empirical date. First we address whether it is possible to find evidence for SLS or the beanbag-selection in actual genomic data. We obtained genome sequence data from 462 fungal species represented in the Fungi division of the ENSEMBL database [26] and calculated the energies for each of the OCCs contained in this database for both the beanbag model and the full model. If the codon distribution was explained by beanbag selection, then we would expect that the energies are distributed multinomially, which would reduce to a binomial distribution in the case of amino acids with 2 codons only. On the other hand, if the genomes are subject to SLS then the full model would be a better description of the data.

To check this, we fitted the frequencies of energies in our dataset to a binomial distribution and to the Boltzmann distribution implied by the full model (i.e. eq. 7). We limited ourselves to the nine amino acids with 2 codons, i.e. 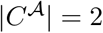, because only for those amino acids it is possible to obtain sufficiently powerful statistics. Furthermore, we only analysed OCCs with lengths 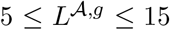. Sequences with *L* < 5 provide too few data points to fit, and for sequences with *L* > 15 the statistical errors become overly large because the number of possible OCCs increases quickly with increasing OCC length. Altogether, we obtained 45702 fits of the fungal data set to the binomial model and the same number of fits to the full model. Similar results with fits to bacterial and protist data are shown in the SI.

### The full model fits the fungal data better than the binomial model

In order to understand how well the genomic data fits the respective models we analysed the mean-residuals of the fits. In the limit of an infinite number of samples that have been generated according to the fitting distribution, the mean-residual would vanish. When the sample size is finite, or when the data is not distributed according to the hypothesis distribution, then the mean-residuals take a positive value. The fits of the fungal data to the binomial distribution yielded mean-residuals between exp(−4) and exp(−9) peaking at about exp(−7); see fig. 2. Visual inspection of a number of examples suggests that these mean-residuals indicate a reasonably good fit to the data.

Next, we fitted the full model eq. 6 to the same data. On the whole, this resulted in smaller mean-residuals, indicating that the full model is a better fit to the data than the binomial model. Fig. 2 summarises this quantitatively. The median for the residuals of the full model is 0.0002850, and as such much smaller than the corresponding value for the binomial fits, which is 0.000845. This suggests that the full model is a better description of the data than the binomial model.

**Fig. 2:**
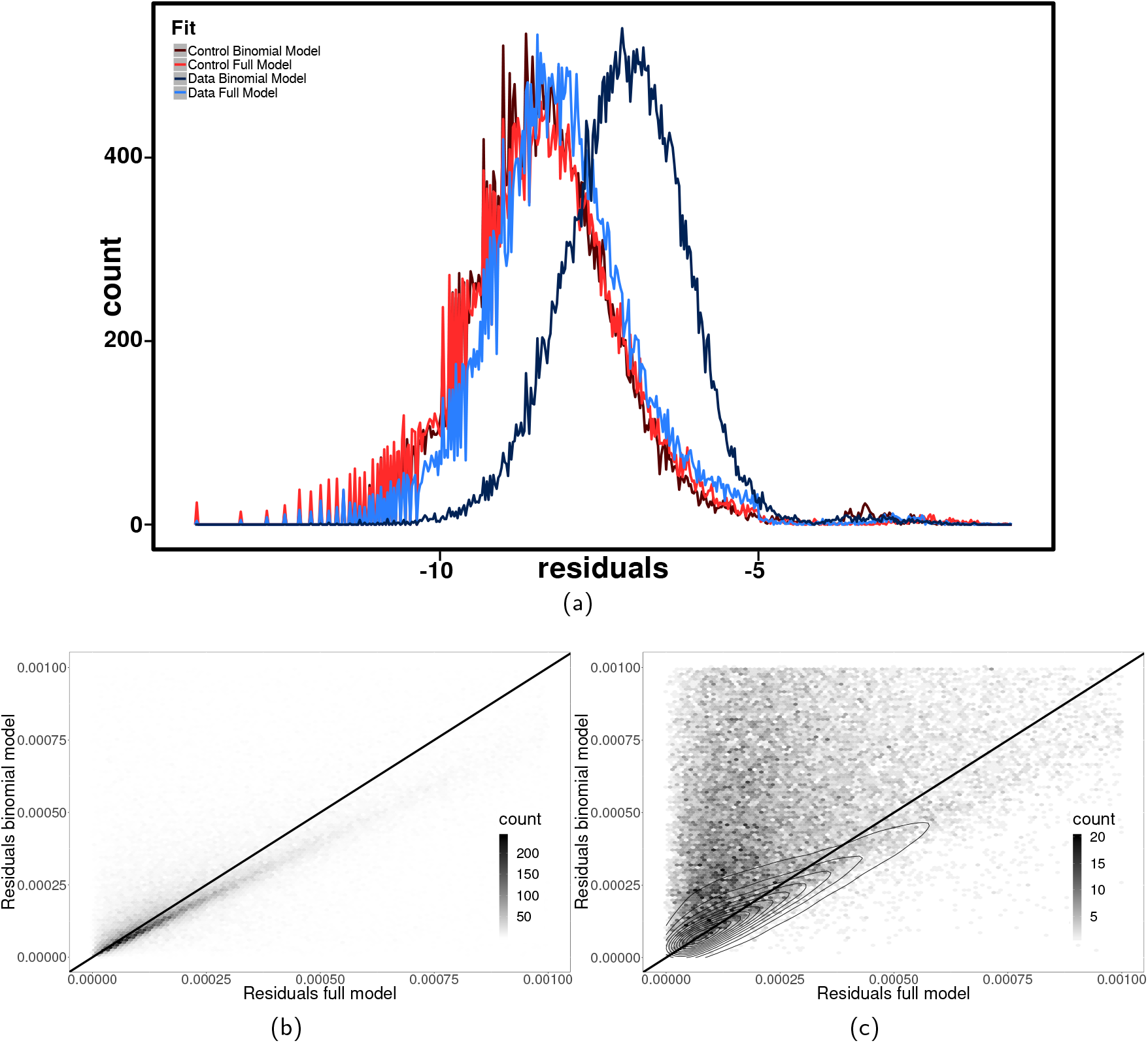
(a) Histogram for the mean-residuals obtained from fitting the binomial distribution and the full model for both the real data and control data generated under the beanbag model assumption, i.e. where codons have been replaced by random synonymous codons with a bias corresponding to the global codon usage bias. The *x*-axis is shown on a logarithmic scale. The distribution of the mean-residuals of the binomial model fitted to real data is clearly shifted to the right relative to the distribution of residual obtained from full model. On the other hand, the mean-residuals of the full model overlap with the distribution of the mean-residuals resulting from the fit of both the binomial and the full model to the control data. (b) Comparing the mean-residuals from the full model to those of the binomial model. The plot shows the density of points for the control data. The area above the diagonal indicates fits where the full model better than the binomial model. Points on the diagonal indicate that both models fit the distribution of OCCs equally well. (c) Same comparison, but for real data. The contour lines indicate the density of the control data in (b) for comparison.

The better fit of the full model could be merely a reflection of the fact that it has more parameters than the binomial model. To check this, we prepared a control set of distributions. This control set consists of the same genomic sequences that the real dataset contains, but with each codon replaced by a random synonymous codon according to the global codon frequency bias *q* of each species; see SI for a description and for the control dataset. By construction, the control set of OCCs implements the beanbag model exactly, i.e. codons are distributed according to a binomial distribution with some species and amino acid dependent probability *q*. Deviations from the binomial distribution in the control set are only due to the statistical error, i.e. a consequence of the finite (and indeed small amount of) data. When we fitted both the full model and the binomial model to the control data, we obtained two distributions of mean-residuals that are visually indistinguishable from one another. This reflects the above cited fact that the full model can approximate control data; see fig. 2a. Interestingly, an inspection of the histogram in fig. 2a reveals that the distribution of mean-residuals obtained from fitting the full model to the real data is only minimally worse (i.e. shifted to the right) compared to the fit to the synthetic data. *Prima facia* this means that almost all the error of the full model fit is due to statistical error, which in turn leads to the conclusion that the full model captures almost all of the variation in the underlying real data.

From the above analyses it is not clear whether the full model is a better fit for all OCCs, or whether it merely fits the majority better while there still are many OCCs that are equally well described by a binomial distribution. In order to investigate this, we compared residuals of fits to the binomial and full models for individual OCCs directly; see figs. 2b & 2c. In order to produce the fits for figure 2c we split gene sequences into OCCs for each of the nine amino acids with a codon choice of 2, and then grouped OCCs by length and amino acid restricting ourselves to the length range 5-15 and the amino acids with 2 codons, as explained above. Thus, any individual organism is represented by up to 9.11 = 99 different residuals, one for each codon and OCC length.

It is instructive to consider the control data first, which by construction follows the binomial distribution. As expected, we found that most OCCs are approximately equally well fitted by the binomial and the full model (see fig. 2b), although the density of points appears to be higher below the diagonal indicating that the binomial model fits the control data somewhat better. This is because, as mentioned above, the full model can only approximate the binomial distribution. In contrast, for the real data the same analysis leads to a high density of points in the upper left corner of the figure, where the mean-residuals of the full model are lower than those of the binomial model; see fig. 2c.

The majority of OCCs are thus better explained by SLS than by beanbag selection. However, there remains a significant minority of OCCs (less than 20%) that can be equally well explained by the beanbag model and SLS. In fig. 2c, these species have points in the *ambiguous region* of the plot (see SI sec. 2 for details on how we defined this region). If we make the assumption that purely by chance some OCCs can be equally well explained by SLS and beanbag selection, then we would expect that all organisms contribute data points to this ambiguous region with equal probability.

Interestingly, the contribution of data points to the ambiguous region is very clearly not governed by chance (fig. 3). Instead, we observe that most organisms completely avoid this region, with the majority of organisms represented with seven data points or fewer (by chance, this should only occur for 0.3% of organisms). On the other hand, a minority of organisms contributes the majority of ambiguous OCC groups, with the highest individual contribution being 51 OCCs (which should occur by chance with a frequency of less than 10^−14^). Thus, while in most organisms SLS is a strong driver of codon usage bias, in some organisms it is weak or absent.

**Fig. 3:**
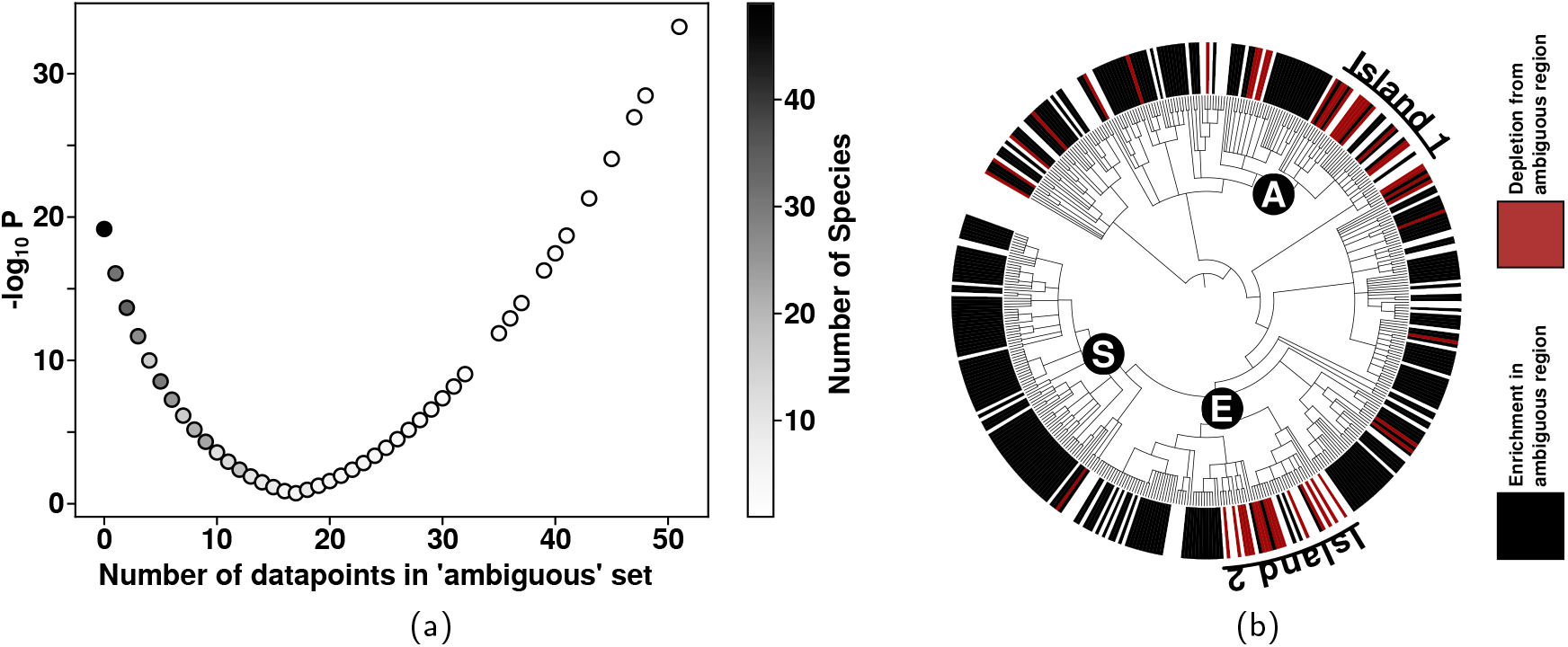
(a) A “Volcano”-plot showing the distribution of points into the ambiguous versus general population classes. A single species could contribute up to 99 data points to the “ambiguous” region highlighted in figure 2c, where it is not possible to distinguish whether SLS or beanbag selection is more likely for a set of OCCs. “ −log_10_ *P*” quantifies the probability of obtaining an observed number of OCCs in the ambiguous region by chance. (b) A taxonomic tree of the species in our fungal dataset, colour coded according to whether the species is statistically under-represented or over-represented in the ambiguous group relative to chance (based on a cut-off value of *P* = 10^−3^). Most species are under-represented indicating that SLS dominates significantly. However, in the two identified “Islands”, for many species SLS does not contribute to shaping codon usage with statistical significance according to our test. Taxonomic groups discussed in the text are indicated (A) *Agaricomycetes;* (E) *Eurotiomycetes*; (S) *Sordariomycetes*.

In the taxonomic tree, organisms where SLS makes apparently weaker contribution to codon usage bias are clearly grouped (fig. 3b). Two larger groups of such organisms are indicated as “Islands” in the figure, and are located within the taxonomic classes *Agaricomycetes* (A) and *Eurotiomycetes* (E). On the other hand, species where SLS makes a weaker contribution to codon usage bias are completely absent from the large *Sordariomycetes* class (S). Thus, absence of clearly detectable SLS appears to be a trait that has evolved in certain taxonomic groups. This is likely linked to the organisms’ lifestyle or other biological traits, although we do not currently understand the exact mechanisms by which the effect of SLS on codon usage could become reduced.

Overall, these observations confirm that codon usage bias is not solely shaped by genomewide forces, but that sequence-level selection makes substantial contributions e.g. via mechanisms linked to translational regulation [9], gene length [27, 28] or other gene specific parameters. Going beyond the current state of the art, we have established here that the distribution of codons can be described compactly by a surprisingly simple distribution, i.e. eq. 6, which can be derived from rather general principle within a statistical physics framework.

### Genome-wide analysis of fungal genomes using the distance measure

For the fungal and bacterial genomes we examined, typical values of the fitted parameters *γ* and *ξ* are small and positive with 96.39% of fits resulting in 0 < *γ*, *ξ* < 2. These fitting parameters do not have an immediate biological interpretation, but interesting insights can be gained by looking in more detail at how species distribute in *ξ*, *γ* space. It is apparent from fig. 4b that in the case of fungal species, the genomic data distributes quite differently than the control data which is based on beanbag selection only. This difference demonstrates that the *global* codon frequency bias q is not the only manifestation of selection pressures on codons, but there are statistical properties to CUB which cannot be captured by a binomial model alone, i.e. there is evidence for SLS.

**Fig. 4:**
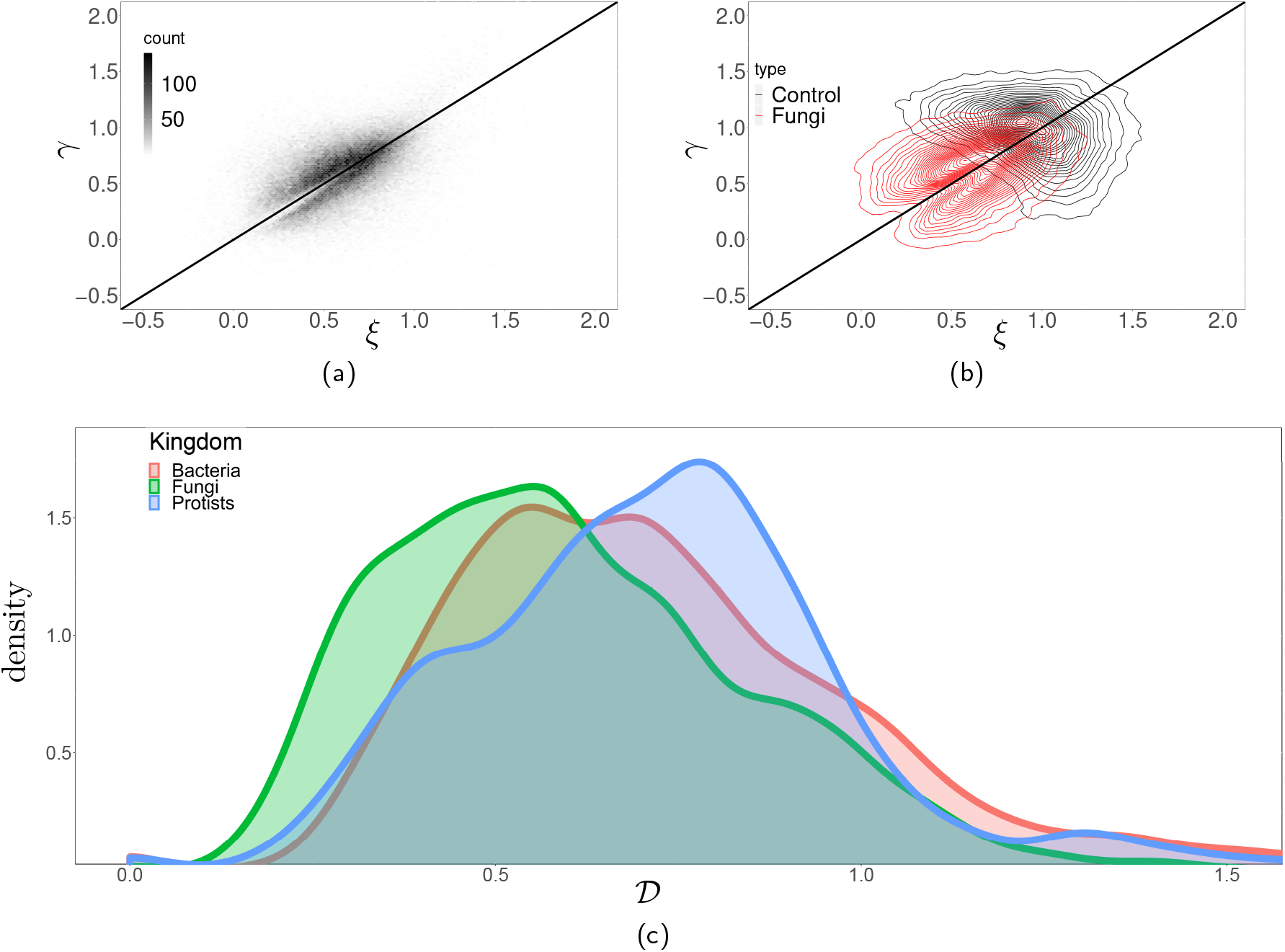
(a) The density of fitted parameters *ξ* and *γ* for each of the 2-codon amino acids for all 462 fungal species in our dataset. We are limiting ourselves to fits with mean-residuals < 0.0009999. The fitted values largely concentrate into the interval of [0, 1.5]. (b) Comparing the fitted parameters obtained from the full model (red) to the fitted parameters obtained from the control (black). The plot shows contour lines of constant density. A comparison with (a) shows that regions of dense points of the control data are restricted to a much smaller area of the parameter space than real genomes. (c) Distribution of the values of 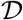 in bacteria, protists and fungi. The plot shows a smoothed distribution of the species averaged distances in the three kingdoms. Clearly, the distances in fungi tends to be the lowest.

We can further probe this using the distance measure 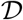 introduced above. 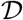 is an organismlevel measure and quantifies how far a genome is from the case of random, unbiased codon usage. It captures two aspects, (*i*) preferential use of one codon over the other and (*ii*) non-random distributions of codons across the genome. In order to gain an intuition for the meaning of 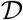, we first compare it to established measures of codon usage bias. Figure 5 displays 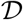 and *F_opt_* (the proportion of optimal codons) as functions of genome complexity. The four analysed genomes are all based on the *S. cerevisiae* genome and contain the same number of sequences encoding identical proteomes. The first genome was generated by replacing all codons with randomly chosen synonymous ones, i.e. this genome has no codon bias at all. For the second genome, codons were again replaced randomly but now with a probability proportional to the observed average codon frequency in the yeast genome: this genome shows the same average codon frequencies as the real genome, but here the bias is generated by a single selective force which applies genome-wide. For the third genome (labelled “Multiple expression dependent biases”), codons were replaced by applying a mixture of two different replacement schemes, with the actual mixture chosen for each gene as a function of its protein expression levels as reported in [29]. This procedure simulates a genome obeying the proposed balance between mutation bias and translation selection underpinning the models described by Gilchrist *et al*., and the parameters chosen for the replacement were selected to recapitulate the data in [9] as closely as possible. The fourth genome is the actual *S. cerevisiae* genome. For these four genomes, we show the distribution of *F_opt_* values for each sequence, as well as amino acid-specific *F_opt_* values averaged over all sequences, and amino acid-specific 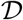-values (note that, because 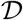 is estimated based on comparing distributions, it is not possible to estimate this for individual sequences). Only values for amino acids encoded by two codons are shown in the latter two cases.

**Fig. 5:**
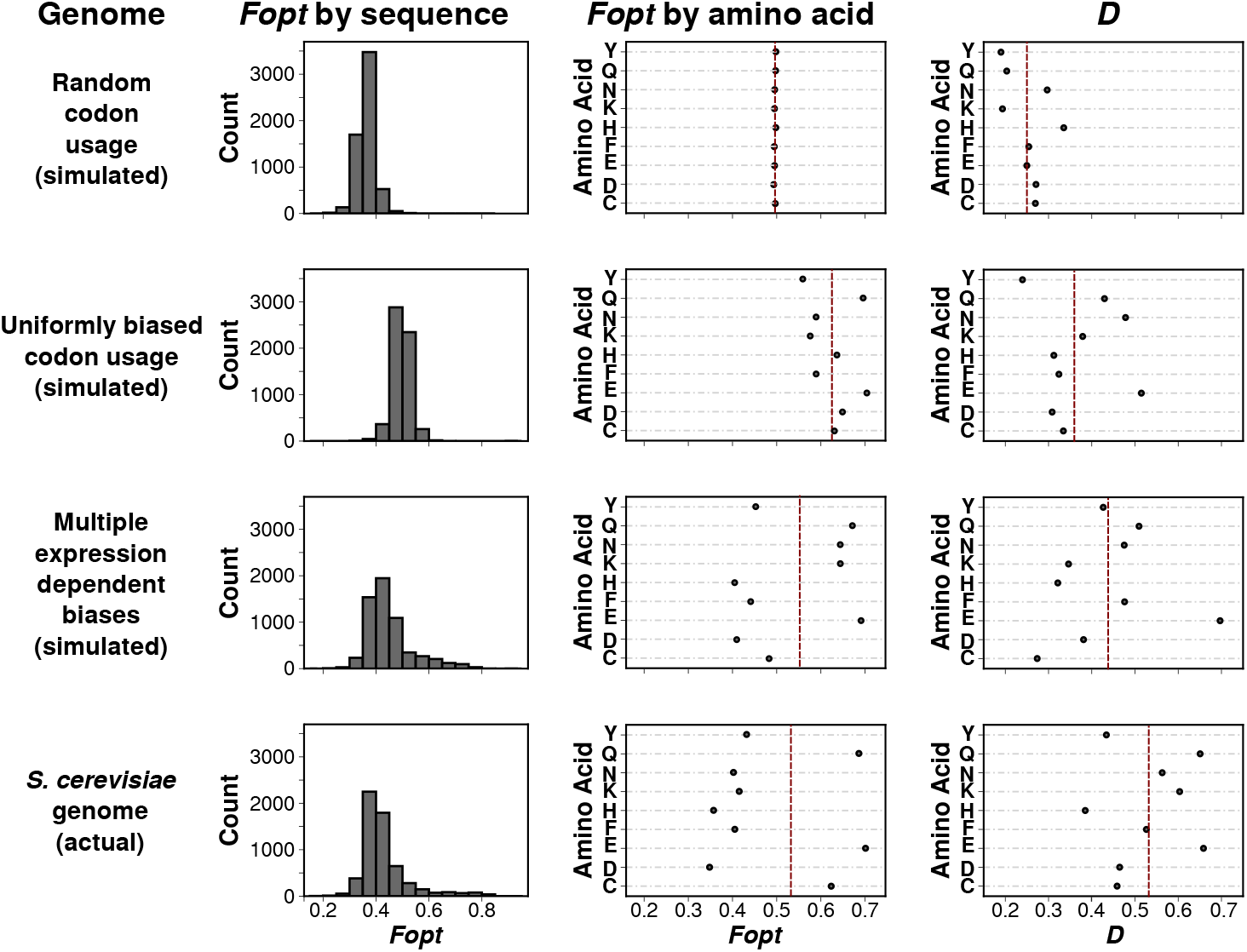
Comparing 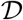 to *F_opt_*. The top three rows analyse simulated genomes encoding identical proteomes as the actual *S. cerevisiae* genome, which is analysed in the bottom row. Columns display *F_opt_* averaged over all amino acids for each sequence (left), *F_opt_* averaged over all sequences for each amino acid with a codon choice 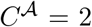 (centre), and 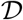 averaged over all sequences for each amino acid with 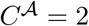 (right). Red lines indicate average values for all analysed amino acids.

A salient property of 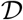 revealed by this analysis is its gradual increase with increasing complexity of the forces that shape codon usage (c.f. the shift of the mean 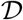 from top to bottom). In contrast, because *F_opt_* simply summarises average sequence properties, its behaviour does not reflect genome complexity and the mean *F_opt_* value shows a non-linear relationship with codon usage bias when complex selective forces shape this bias. Although CAI and *CBI* are calculated differently, they correlate with *F_opt_* and would show a similar, non-linear relationship. Interestingly, there is a notable increase in 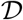 between the complex simulated and the actual yeast genome, indicating that the actual yeast genome is shaped by a more complex array of forces than the mutation bias/translational selection model alone accounts for.

One motivation for proposing 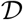 as a useful measure for characterising codon usage in fungal genomes was the fact that this measure does not require any context information other than the nature of the coding sequences themselves, and is thus applicable to any genome irrespective of the degree of knowledge available on the corresponding organism. We calculated the average 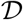 for each of the many hundreds of species with genome information in the ENSEMBL Fungi database [26], as well as for a size-matched selection of bacterial genomes. Fig. 6 shows the resulting 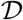 values, grouped by taxonomic class of the species. This analysis highlights a number of features that are not apparent from existing analyses. Within each taxonomic class there is significant diversity in 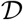 values, indicating that the diversity and nature of forces that shape codon usage varies widely even within taxonomic classes. This trend is particularly noticeable within the fungi whereas most bacterial classes display more uniform 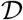 values (although there are exceptions, e.g. within the *Mollicutes* and *Deinococci*). Despite the high degree of variability within classes, 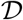 clearly arises from heritable traits, since the average difference in 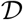 between two species increases proportionally with the taxonomic distance between these species (fig. 6 B,D). Interestingly, the relationship between 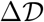 and taxonomic distance is quantitatively different between fungi and bacteria (compare the slopes in the inset graphs in Fig. 6.)

**Fig. 6:**
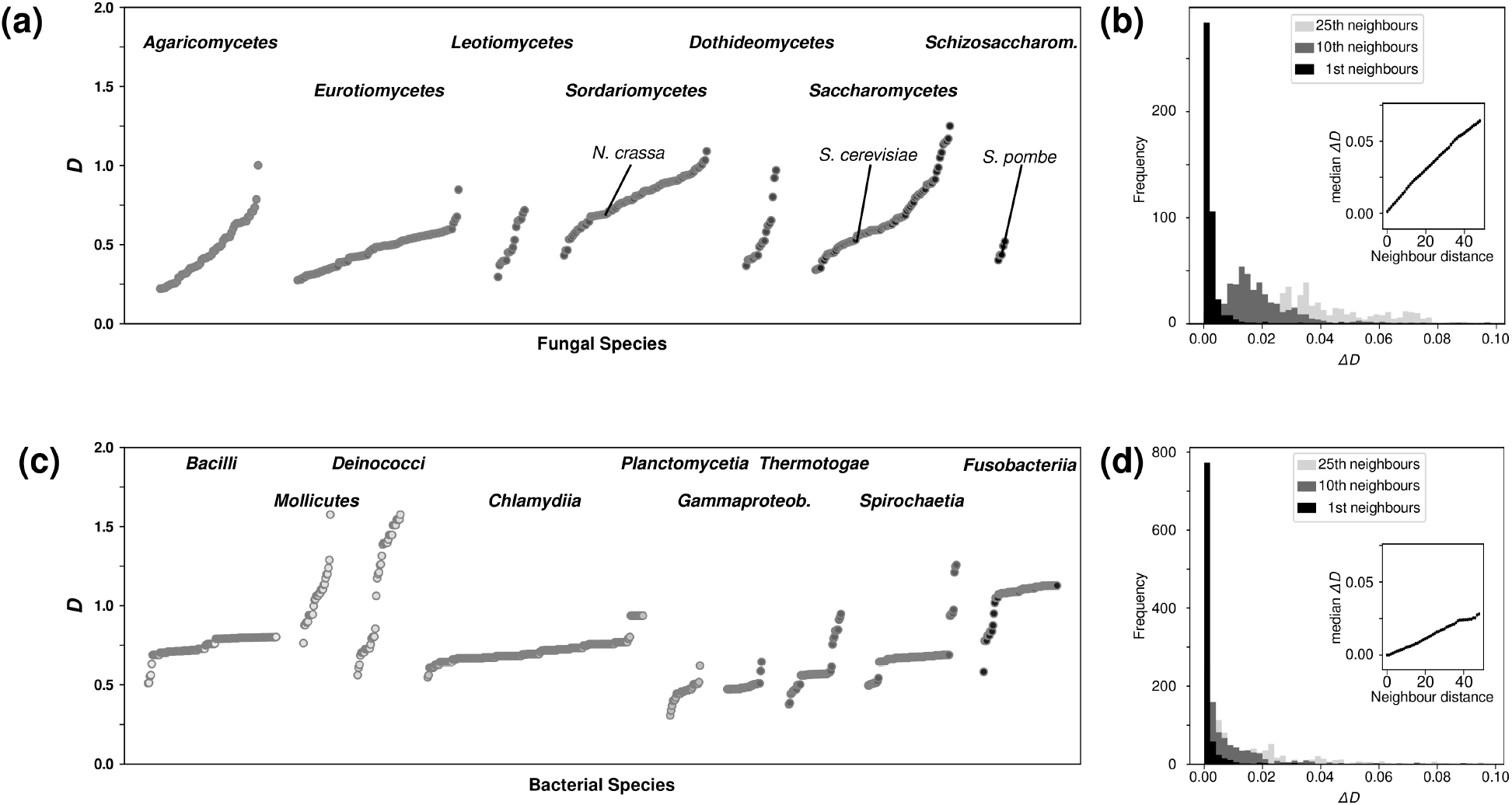
Genome-wide 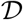 for fungal (top) and selected bacterial (bottom) species. A, Genome-wide 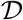 for selected fungal taxonomic classes with multiple sequenced genomes. B, the difference in 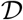 between species increases with taxonomic distance. Detailed histograms show values for 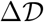 for species pairs that are immediate taxonomic neighbours (black),or that are separated by 10 (dark grey) or 25 (light grey) species. Inset, median 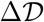 as a function of neighbour distance. C and D, as in A and B but for bacterial species.

As an additional analysis, we asked whether organismal lifestyle can significantly influence 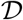. We analysed the descriptive paragraphs accompanying genome sequencing projects on the ENSEMBL Fungi database for indicators of specific lifestyles, including whether individual species were known pathogens or symbionts of other organisms. Of the lifestyles assessed in this analysis, we observed significantly lower 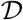 values for mycorrhizal and brown rot fungi, with very strong significance (*p* < 10^−6^) for the former (figure 7). Mycorrhizal fungi are a phylogenetically diverse group that live in symbiotic relationships with plant roots, with which they exchange nutrients. It has been suggested that associations between mycorrhizal fungi and plants are unstable in evolutionary terms, with frequent reversals to free-living conditions [30]. If this was indeed the case, one explanation for the low 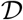 values in these organisms could be that their genomes are not in steady state, and that eq. 7 thus may not capture their genome dynamics accurately. Interestingly, the other group of symbiotic fungi contained in this analysis, endophytic fungi, are not associated with particularly low 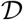 values, indicating that this is not a general consequence of symbiotic lifestyles. Overall however, these analyses confirm that particular ecological niches *can* be associated with particular 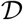 values, although this is not a general rule.

**Fig. 7:**
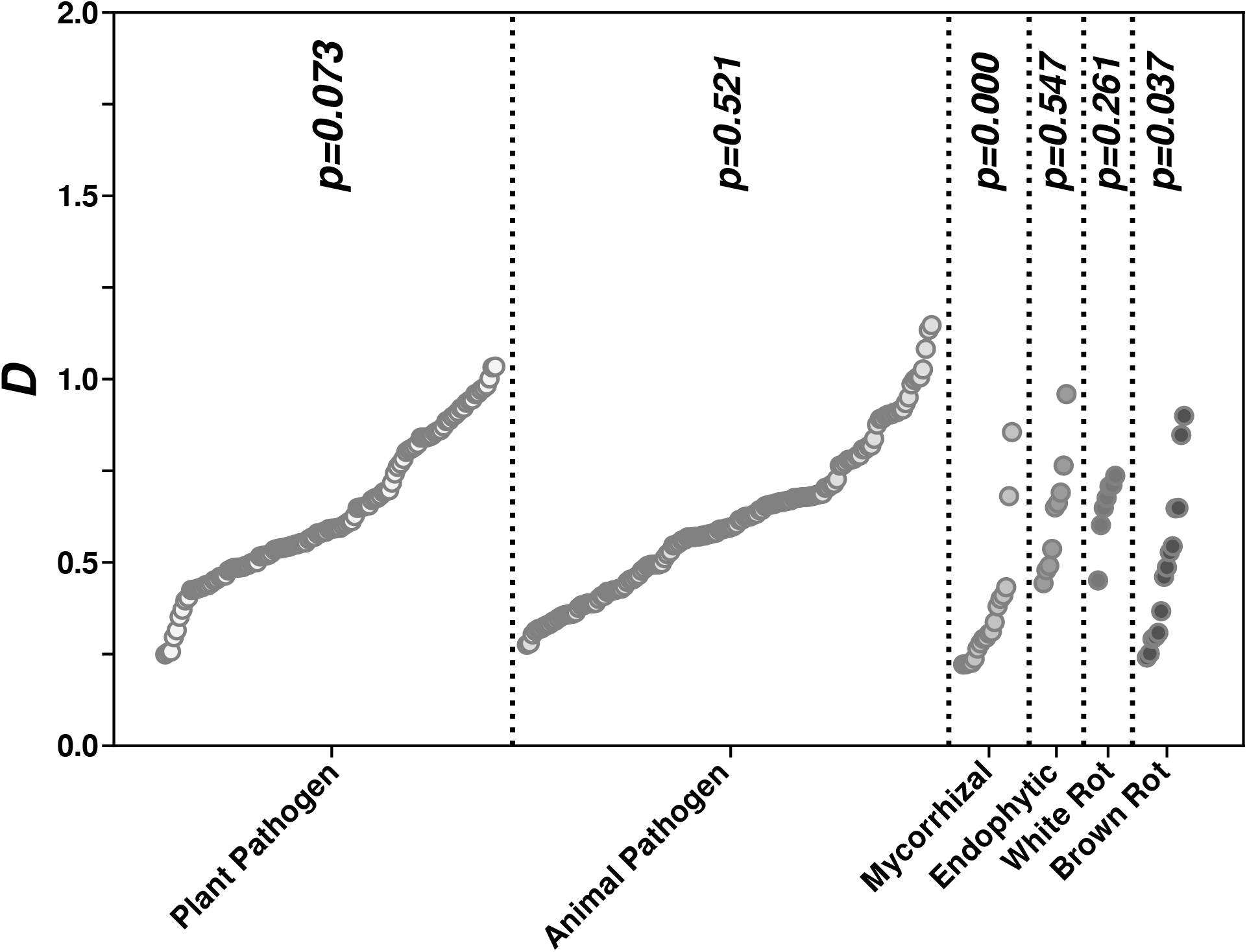
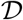 values in fungi with distinct lifestyles. *p*-values indicate significantly different distribution of 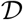 in a particular lifestyle compared to the full 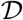 distribution. Significance of the difference in distribution of 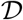 between individual lifestyles and fungi overall was assessed using a non-parametric Kruskal-Wallis test.

### 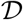 reveals amino acid-specific patterns of codon selection pressure

An advantage of using 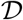 over other measures of codon usage bias is that it lends itself to detecting differences in selective pressure in different OCC sets. By way of example, we compared how 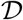 differs for different amino acids in the fungal kingdom. Initial visual inspection of the dataset revealed that, as a general pattern, most amino acids in the same organism behave similarly in terms of 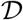, suggesting that they experience similar selective forces. There are, however, also exceptions to this pattern. Fig. 8 reveals that atypically stronger or weaker selection for particular amino acids is an evolutionary feature that is linked to taxonomic groups. In this analysis, we define atypical selection as a 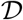 value that is more than 2 standard deviations above or below the average 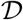 value for that organism. Particularly notable patterns include the *Sordariomycetes* group where the amino acid phenylalanine (F) shows atypically strong codon selection (higher than average 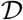 values) in most species, whereas in other groups F either behaves like other amino acids or shows atypically weak selection (eg in the *Agaricomycetes*). The fact that codon bias differs for different amino acids in any one organism has been observed before and is both apparent from inspecting average codon usage tables, and predicted from current models of codon usage bias evolution such as ref. [9]. A novel insight emerging from our analyses is how strongly the direction of these differences can vary between taxonomic groups. Because codons are decoded by tRNAs, an immediate hypothesis could be that atypical 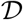 values reflect unusual features of the tRNA population. This would make sense as tRNA levels are well known to affect codon decoding speed, ribosome movement, and thereby translational control [31]. Moreover, a detailed inspection of figure 5 reveals that the three amino acids encoded by A/G ending codons, which are decoded by two separate tRNAs, have higher 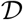 values than the six amino acids encoded by C/U ending codons which are decoded by a single, wobbling tRNA (cf row “*S. cerevisiae*genome (actual)”, column 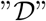 in this figure). While we cannot mechanistically explain this observation, it does suggest that at least some aspects of 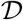 can be explained by the physical nature of the tRNA decoding system.

**Fig. 8:**
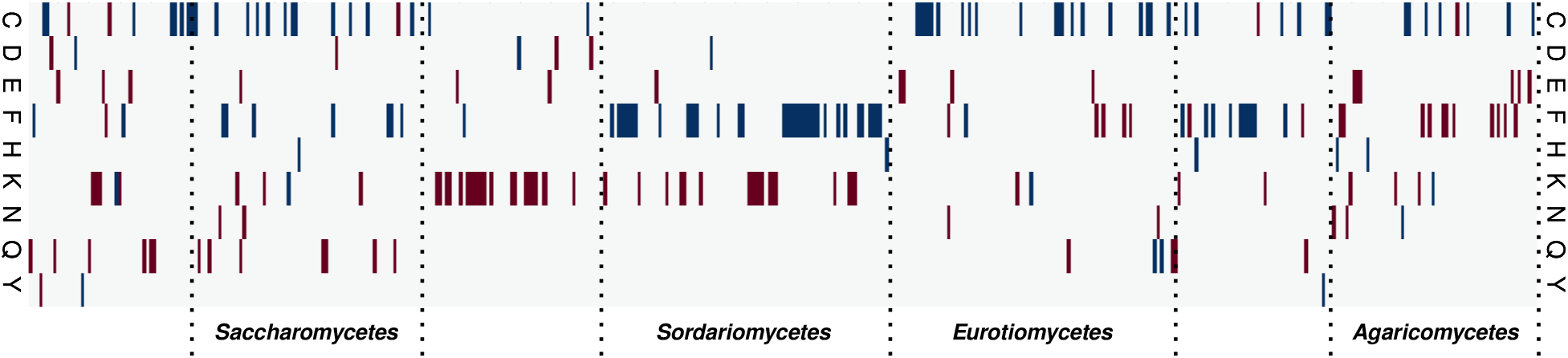
Amino acid-specific patterns of codon usage bias in fungal genomes. Average 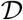 values were calculated for all each amino acid in each genome. Amino acids are highlighted if their 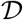 value was more than 2*σ* above (blue) or below (red) the median 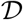 for that species, i.e. if they are under atypical selection compared to other amino acids in the same species. Species were ordered according to the taxonomic hierarchy in NCBI taxonomy, and taxonomic groups represented with larger numbers of genomes are indicated.

To explore this issue further, we identified tRNA genes in all fungal genomes included in our study using tRNAscan-SE software [32]. We then used a regression approach to explore in how far tRNA gene copy number could explain atypical 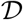 values for particular amino acids. The lasso regression approach chosen for this analysis [33] allows estimating both the predictive power of a dataset for particular target variables, and the importance of individual dataset features for the prediction.

The results of this analysis show that overall, the tRNA pool has variable predictive power over amino acid specific 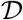 in individual organisms (Fig. 9). The A/G ending amino acids Q and K are again conspicuous as the two amino acids for which tRNA gene copy number has the greatest explanatory power over D values, although for the third A/G ending amino acid, E, 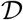 is less well explained by the tRNA pool. Although for Q and K tRNAs explains 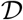 to *some extent*, we note that even for these amino acids the absolute explanatory power of the tRNA pool appears remarkably low, and for other amino acids like H tRNA gene copy number cannot explain 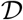 at all. Thus, tRNA abundance appears as only one contributory factor among potentially many that shape codon usage patterns consistent with our initial assumption of many contributory forces shaping CUB.

**Fig. 9:**
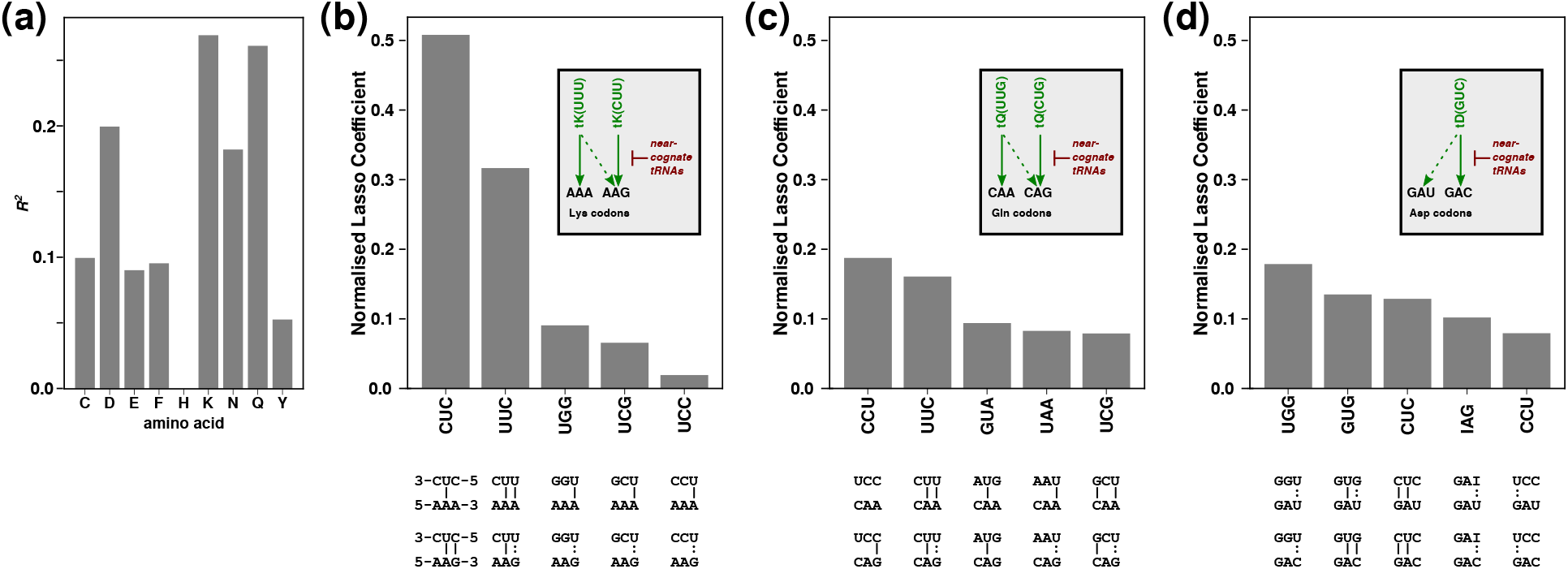
tRNA gene copy number can only partially explain atypical 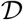 values. A, Lasso regression-derived *R*^2^ values for tRNA gene copy numbers and 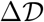 values (the difference between 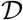 for an individual amino acid and the average 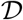 for all amino acids in the same organism). *R*^2^ is proportional to the predictive power of tRNA gene copy number to explain atypical 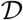. B-D, importance of individual tRNAs for predicting atypical 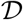 for the three amino acids where tRNA gene copy number has the strongest predictive power over 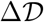. The five strongest predicting tRNAs are shown for each amino acid, and their base-pairing potential with the codons encoding the amino acid in question is indicated below the main panels (|, Watson Crick base-pair;:, wobble base-pair). The inset figures illustrate the predominant decoding schemes for each amino acid in fungi.

For the three amino acids where 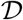 is best explained by the tRNA pool, we recovered the individual Lasso coefficients which are indicators of the contribution the corresponding tRNA makes to the predictive power of the pool. In the case of lysine (K), more than 80% of the explanatory power lies with the gene copy number of two glutamic acid inserting tRNAs, tE(CUC) and tE(UUC) (Fig. 9 B). Both tRNAs can form two Watson-Crick base pairs with one or other of the lysine codons, which makes them strong candidates to be near-cognate tRNAs for these codons (ie tRNAs which inhibit efficient decoding, [31, 34]). For glutamine (Q) and aspartic acid (D), the power to explain 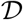 is spread out over many more tRNAs than is the case for K, and here the top two tRNA species contribute less than 40 % (Fig. 9 C,D). In both cases however, the top five tRNAs include several which can form Watson-Crick or wobble base pairs with the codons, making them potential candidates as near-cognate tRNAs. Since the top scoring tRNAs for both Q and D do not strongly interact with the codons in questions however, near-cognate interactions with codons are certainly not the only mechanism by which the tRNA pool shapes codon usage patterns of genomes.

In sum, these data indicate that in some of the cases where atypical selection is observed forindividual amino acids, this is in part in response to the particular tRNA pools in these organisms. However, currently uncharacterised influences are also at play, and for some amino acids atypical codon selection is entirely caused by forces unrelated to tRNA abundance.

## Discussion

The model we propose draws heavily upon ideas from statistical mechanics and especially stochastic thermodynamics [24, 35]. Its only assumption is that genomes are the result of a random walk and that they are in steady state. This is in contrast to previous models of codon-specific evolutionary selection that typically assume a small number of specific drivers of selection, compared to the many potential drivers that have been proposed in the literature. This agnosticism with respect to the origins of CUB makes our model also more easily applicable. For example, it does not require an a *priori* reference set.

The model led directly to a novel measure of codon usage bias, i.e. the distance 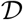 that encapsulates both the global codon usage bias due to global frequency imbalances (≈ bias due to beanbag model) and the codon usage bias resulting from non-random distributions of codons across genes (≈ bias due to SLS).

While 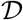 captures two aspects of CUB, a more detailed analysis makes it possible to disentangle these. From fig. 4a it is clear that the distribution of genomes in *ξ*, *γ* space is very different from the distribution of hypothetical genomes that are only subject to selection by the beanbag model. Indeed, there is a sharp demarcating line in this space across which beanbag model genomes cannot be found.

While 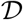 has the ability to explain codon selection in greater depth and does not require any contextual datasets or assumptions it is not suitable to analyse individual genes, or even small sets of genes. This is because it is based on fitting a model to a distribution of OCCs. This can only be done if the underlying data yields a sufficiently accurate statistics.

Our application of 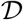 to fungal genomes represented in the fungal section of the ENSEMBL database [26], as well as a size-matched selection of genomes from the bacterial section of the same database yielded interesting results. These show that the nature and diversity of selective forces acting on codon usage is surprisingly varied even in taxonomic classes within these groups, as indicated by the spread in 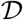 values (Fig. 6), even though 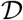 is clearly based on heritable traits. The fact that common model organisms tend to be located near the median of the range underlines that it may not be appropriate to simply port approaches for analysis of codon usage from such models to other organisms. For example, while models relying on the analysis of opposing forces of mutation bias and translational selection [9] appear to come relatively close to describing codon usage bias in baker’s yeast 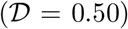, the diversity of forces acting on codon usage appears very different in other fungi of interest, including human pathogens like *Candida albicans* 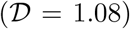, *Candida glabrata* 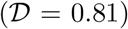 or *Pneumocystis carinii* 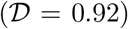; and plant pests like *Verticillum* 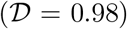 and *Magnaporthe* 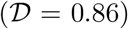.

### The dataset

All datasets were obtained from ENSEMBL https://www.ensembl.org. We downloaded coding sequences for 462 species from the fungal kingdom (release 36 in August 2017), and 442 randomly chosen species from the bacterial kingdom (release 40 in July 2018). Bacterial species were chosen to roughly size-match the fungal data set, while proportionally representing the taxonomic spread of sequenced bacterial genomes. All species names and corresponding download weblinks are in supplementary file “species.xlsx”. We then produced clean sequence files by converting each gene sequence into a valid codon sequence and removing those genes where the number of nucleotides was not a multiple of 3 (indicating errors in the ORF annotation). This led to the exclusion of 35748 genes from 4554328 total genes in the fungal dataset, and of 6384 genes from 1286467 in the bacterial dataset.

## References

[1] Ikemura T. Correlation between the abundance of Escherichia coli transfer RNAs and the occurrence of the respective codons in its protein genes: a proposal for a synonymous codon choice that is optimal for the E. coli translational system. Journal of Molecular Biology. 1981;151(3):389–409.

[2] Bennetzens JL, Hall BD. Codon Selection in Yeast. The Journal of Biological Chemistry. 1982;257(6):3026–3031.

[3] Sharp PM, Li WH. The codon Adaptation Index–a measure of directional synonymous codon usage bias, and its potential applications. Nucleic Acids Research. 1987;15(3):1281–1295.

[4] Wright F. The ‘effective number of codons’ used in a gene. Gene. 1990;87(1):23–29.

[5] Shah P, Gilchrist MA. Explaining complex codon usage patterns with selection for translational efficiency, mutation bias, and genetic drift. Proceedings of the National Academy of Sciences of the United States of America. 2011;108(25):10231–10236. doi:10.1073/pnas.1016719108.

[6] Gilchrist MA. Combining Models of Protein Translation and Population Genetics to Predict Protein Production Rates from Codon Usage Patterns. Molecular Biology and Evolution. 2007;24(11):2362–2372. doi:10.1093/molbev/msm169.

[7] Sharp PM, Bailes E, Grocock RJ, Peden JF, Sockett RE. Variation in the strength of selected codon usage bias among bacteria. Nucleic Acids Research. 2005;33(4):1141–1153. doi:10.1093/nar/gki242.

[8] Ran W, Kristensen DM, Koonin EV. Coupling Between Protein Level Selection and Codon Usage Optimization in the Evolution of Bacteria and Archaea. mBio. 2014;5(2):e00956. doi:10.1128/mBio.00956-14.

[9] Gilchrist MA, Chen WC, Shah P, Landerer CL, Zaretzki R. Estimating Gene Expression and Codon-Specific Translational Efficiencies, Mutation Biases, and Selection Coefficients from Genomic Data Alone. Genome Biology and Evolution. 2015;7(6):1559–1579. doi:10.1093/gbe/evv087.

[10] Bulmer M. The selection-mutation-drift theory of synonymous codon usage. Genetics. 1991;129(3):897–907.

[11] Chu D, von der Haar T. The architecture of eukaryotic translation. Nucleic Acids Research. 2012;40(20):10098–10106. doi:10.1093/nar/gks825.

[12] Chu D, Kazana E, Bellanger N, Singh T, Tuite M, von der Haar T. Translation elongation can control translation initiation on eukaryotic mRNAs. EMBO Journal. 2014;33(1):21–34. doi:10.1002/embj.201385651.

[13] Neafsey D, Galagan J. Positive selection for unpreferred codon usage in eukaryotic genomes. BMC Evolutionary Biology. 2007;7(1):119. doi:10.1186/1471-2148-7-119.

[14] Zhou T, Weems M, Wilke C. Translationally Optimal Codons Associate with Structurally Sensitive Sites in Proteins. Molecular Biology and Evolution. 2009;26(7):1571–1580. doi:10.1093/molbev/msp070.

[15] Biro JC. Does codon bias have an evolutionary origin? Theoretical Biology and Medical Modelling. 2008;5(1):16. doi:10.1186/1742-4682-5-16.

[16] Trotta E. Selection on codon bias in yeast: a transcriptional hypothesis. Nucleic Acids Research. 2013;41(20):9382–9395. doi:10.1093/nar/gkt740.

[17] Yannai A, Katz S, Hershberg R. The Codon Usage of Lowly Expressed Genes Is Subject to Natural Selection. Genome Biology and Evolution. 2018;10(5):1237–1246. doi:10.1093/gbe/evy084.

[18] Southworth J, Armitage P, Fallon B, Dawson H, Bryk J, Carr M. Patterns of Ancestral Animal Codon Usage Bias Revealed through Holozoan Protists. Molecular Biology and Evolution. 2018;35(10):2499–2511. doi:10.1093/molbev/msy157.

[19] Zhou Z, Dang Y, Zhou M, Yuan H, Liu Y. Codon usage biases co-evolve with transcription termination machinery to suppress premature cleavage and polyadenylation. eLife. 2018;7. doi:10.7554/eLife.33569.

[20] Jossé L, Singh T, von der Haar T. Experimental determination of codon usage-dependent selective pressure on high copy-number genes in Saccharomyces cerevisiae. bioRxiv. 2018; doi:10.1101/358259.

[21] Xiang H, Zhang R, Butler RR, Liu T, Zhang L, Pombert JF, et al. Comparative Analysis of Codon Usage Bias Patterns in Microsporidian Genomes. PLOS ONE. 2015;10(6):e0129223. doi:10.1371/journal.pone.0129223.

[22] Badet T, Peyraud R, Mbengue M, Navaud O, Derbyshire M, Oliver RP, et al. Codon optimization underpins generalist parasitism in fungi. eLife. 2017;6. doi:10.7554/eLife.22472.

[23] Sinha I, Woodrow CJ. Forces acting on codon bias in malaria parasites. Scientific Reports. 2018;8(1):15984. doi:10.1038/s41598-018-34404-9.

[24] Seifert U. Stochastic thermodynamics, fluctuation theorems and molecular machines. Reports on Progress in Physics. 2012;75(12):126001.

[25] Crooks G. Nonequilibrium Measurements of Free Energy Differences for Microscopically Reversible Markovian Systems. Journal of Statistical Physics. 1998;90(5/6):1481–1487. doi:10.1023/a:1023208217925.

[26] Kersey PJ, Allen JE, Allot A, Barba M, Boddu S, Bolt BJ, et al. Ensembl Genomes 2018: an integrated omics infrastructure for non-vertebrate species. Nucleic Acids Res. 2018;46(D1):D802–D808.

[27] Duret L, Mouchiroud D. Expression pattern and, surprisingly, gene length shape codon usage in Caenorhabditis, Drosophila, and Arabidopsis. Proc Natl Acad Sci USA. 1999;96(8):4482–4487.

[28] Moriyama EN, Powell JR. Gene length and codon usage bias in Drosophila melanogaster, Saccharomyces cerevisiae and Escherichia coli. Nucleic Acids Res. 1998;26(13):3188–3193.

[29] Ho B, Baryshnikova A, Brown GW. Unification of Protein Abundance Datasets Yields a Quantitative Saccharomyces cerevisiae Proteome. Cell Systems. 2018;6(2):192–205.e3. doi:10.1016/j.cels.2017.12.004.

[30] Hibbett D, Gilbert L, Donoghue M. Evolutionary instability of ectomycorrhizal symbioses in basidiomycetes. Nature. 2000;407(6803):506–508. doi:10.1038/35035065.

[31] Tarrant D, von der Haar T. Synonymous codons, ribosome speed, and eukaryotic gene expression regulation. Cellular and Molecular Life Sciences. 2014;71(21):4195–4206.

[32] Lowe TM, Eddy SR. tRNAscan-SE: A Program for Improved Detection of Transfer RNA Genes in Genomic Sequence. Nucleic Acids Research. 1997;25(5):955–964. doi:10.1093/nar/25.5.955.

[33] Tibshirani R. Regression Shrinkage and Selection Via the Lasso. Journal of the Royal Statistical Society: Series B (Methodological). 1996;58(1):267–288. doi:10.1111/j.2517-6161.1996.tb02080.x.

[34] Plant EP, Nguyen P, Russ JR, Pittman YR, Nguyen T, Quesinberry JT, et al. Differentiating between near- and non-cognate codons in Saccharomyces cerevisiae. PLoS ONE. 2007;2(6):e517.

[35] Parrondo J, Horowitz J, Sagawa T. Thermodynamics of information. Nature Physics. 2015;11(2):131–139. doi:10.1038/nphys3230.

